# Modulation of neural activity in frontopolar cortex drives reward-based motor learning

**DOI:** 10.1101/2020.05.05.077743

**Authors:** M Herrojo Ruiz, T Maudrich, B Kalloch, D Sammler, R Kenville, A Villringer, B Sehm, V Nikulin

## Abstract

Decision-making is increasingly being recognised to play a role in learning motor skills. Understanding the neural processes regulating motor decision-making is therefore essential to identify mechanisms that contribute to motor skill learning. In decision-making tasks, the frontopolar cortex (FPC) is involved in tracking the reward of different alternative choices, as well as their reliability. Whether this FPC function extends to reward landscapes associated with a continuous movement dimension remains unknown. Here we used anodal transcranial direct current stimulation (tDCS) over the right FPC to investigate its role in reward-based motor learning. Nineteen healthy human participants completed a motor sequence learning task using trial-wise reward feedback to discover a hidden performance goal along a continuous dimension: timing. As a control condition, we modulated contralateral motor cortex (left M1) activity with tDCS, which has been shown to benefit motor skill learning but less consistently reward-based motor learning. Each active tDCS condition was contrasted to sham stimulation. Right FPC-tDCS led to faster learning primarily through a regulation of exploration, without concurrent modulation of motor noise. A Bayesian computational model revealed that following rFPC-tDCS, participants had a higher expectation of reward, consistent with their faster learning. These higher reward estimates were inferred to be less volatile, and thus participants under rFPC-tDCS deemed the mapping between movement and reward to be more stable. Relative to sham, lM1-tDCS did not significantly modulate main behavioral outcomes. The results indicate that brain regions previously linked to decision-making, such as the FPC, are relevant for motor skill learning.

## Introduction

One of the hallmarks of motor skill learning is the reduction in movement variability [1, 2]. As the dancer learns to perform pirouettes, the initial irregularity in the movement amplitude and timing decreases and the turns become smoother. In this context, movement variability is regarded as motor noise, and is dissociated from the intentional use of motor variability, termed motor exploration [3, 4]. While learning motor skills leads to improvements in movement execution, it also influences action selection [5]. For instance, an expert musician learns to decide upon the most suitable phrasing for a piece and style. Thus, motor skill learning can be understood as a decision-making process, a view shared by an increasing number of studies [6, 4, 7, 8]. In this framework, motor decision-making involves an initial exploration of movement alternatives in a continuous space, which is followed by the exploitation of inferred optimal movements and gradually refined by reducing motor noise [9]. But if motor skill learning depends on making the right decisions about a movement, a fitting prediction would be that brain regions involved in regulating the exploration-exploitation tradeoff in cognitive decision-making tasks would also modulate motor decision making.

Here, we postulate that the most anterior part of the human prefrontal cortex, the frontopolar cortex (FPC), may have a crucial role in driving motor decision-making when external reward signals are available to drive exploration-exploitation. In decision-making tasks involving two or more choices, the right FPC has been identified as promoting directed exploration, driven by information seeking and extrinsic feedback [10, 11]. Directed exploration in this context is conveived as independent from random exploration, also related to the intentional use of variability but in the absence of external feedback guiding the process. In addition, FPC has been linked to tracking the reward value associated with different options, strategies or goals [12] and with the rapid learning of novel rules [13]. Beyond the context of decision-making tasks, it is reasonable to assume that FPC would also influence the exploration-exploitation balance during motor learning. The rationale is that motor learning takes place in a continuous movement space [6]; the capability of FPC to monitor multiple discrete choices and their reward would make it an ideal candidate to track the reward associated with continuous movement parameters.

To identify the role of the FPC in motor decision-making, we used noninvasive brain stimulation (NIBS). NIBS techniques, such as transcranial magnetic stimulation (TMS) or transcranial direct current stimulation (tDCS), have been extensively used to modulate motor performance or motor learning in different scenarios [14, 15, 16, 17]. So far, these stimulation techniques have primarily targeted cortical motor brain regions, such as the primary motor cortex (M1) or supplementary motor area (SMA). In tDCS studies, there is evidence that changes in motor cortical excitability induced by M1 or SMA tDCS can have a positive effect on different aspects of motor learning, including explicit or implicit sequence learning and visuomotor-adaptation tasks [18, 15, 19, 20]. Reward-based motor learning, however, had not been the focus of tDCS investigations until recently [28]. This study demonstrated that combining reward signals with M1-tDCS could improve motor learning, as reflected in the retention of motor skills. More consistent evidence for a role of M1 in processing reward signals during motor learning comes from TMS studies. Specifically, using M1-TMS to assess motor-evoked potentials, different authors have demonstrated that motor cortex excitability is enhanced in anticipation of rewarding movements [21, 22], and more generally, upcoming appetitive stimuli [23, 24]. However, it is still unclear whether M1-tDCS consistently affects reward-based motor learning and, thus, motor decision-making. In addition, the effects of NIBS outside the motor system on the modulation of motor learning remain elusive.

To probe the functional relevance of the FPC in regulating exploration and controlling the intentional use of motor variability during reward-based motor learning, we used anodal tDCS over the right FPC. We selected the right over the left FPC due to its greater engagement in regulating the exploration-exploitation balance [25, 11]. Nineteen healthy participants completed a motor sequence learning task using trial-wise reward feedback to discover a hidden performance goal: a timing pattern. It has been argued that in the real world the acquisition of complex motor skills often involves an uncertain or variable mapping between actions and outcomes [26]. Moreover, higher reward uncertainty can be beneficial for motor retention [26]. Here we also implemented a mapping rule between movement and reward governed by uncertainty, as in our task different timing patterns could receive the same reward, whereas similar timing patterns would obtain different rewards. Accordingly, higher overall exploration would lead to the perception of higher uncertainty in the environment (or greater estimated change in the reward tendency, termed environmental volatility, [27]). In this scenario, the optimal strategy to maximize the mean total reward would be to keep track of the action-reward associations during initial exploration, and to swiftly switch to exploitation of the performance inferred as most rewarding. As control tDCS condition we modulated contralateral motor cortex activity using contralateral (left) M1 tDCS —shown to positively influence different components of motor skill learning with a less established modulation of reward processing [19, 20, 28]. Each active tDCS condition was contrasted to sham stimulation. The effect of the tDCS stimulation protocols on the individual brain was further assessed using simulations of the electric field strength guided by individual T1-weighted anatomical magnetic resonance images (MRI).

The central hypothesis was that rFPC-tDCS modulates the exploration-exploitation balance during reward-based motor learning to increase the total reward relative to sham. Additionally, we predicted that the rFPC-tDCS effects would be limited to the regulation of the intentional use of motor variability —motor exploration —without concurrent changes in motor noise. By contrast, lM1-tDCS was expected to affect motor noise but not motor exploration when compared to sham stimulation. Motor noise represents the residual variability that is expressed when aiming to accurately reproduce the same action [29, 4]. To measure motor exploration, we took an independent measurement of each participant’s motor noise and subtracted it from the total level of motor variability expressed during reward-based motor learning [4].

## Results

Nineteen participants took part in our double-blind longitudinal study on the effects of rFPC-tDCS on reward-based motor learning. Participants underwent each of three types of a tDCS protocol (rFPC, lM1, sham condition) on separate days over three weeks in a pseudo-randomized counterbalanced order across participants. Selection of target coordinates for each tDCS protocol was guided by individual T1-MRI. In each tDCS session, participants completed our reward-based motor sequence learning task with their right hand on a digital piano (Yamaha Clavinova CLP-150, Yamaha Corporation, Hamamatsu, Japan). The task consisted of an initial baseline phase of 20 trials of regular isochronous performance, followed by three blocks of 30 trials each of reward-based learning (Figure 1A). The baseline phase required participants to press a series of eight consecutive white piano keys with four fingers (four notes upwards + same four notes downwards; one finger per key with fixed finger-to-key mapping) regularly at a self-paced tempo. This phase allowed us to assess baseline motor noise during regular performance.

**Figure 1:**
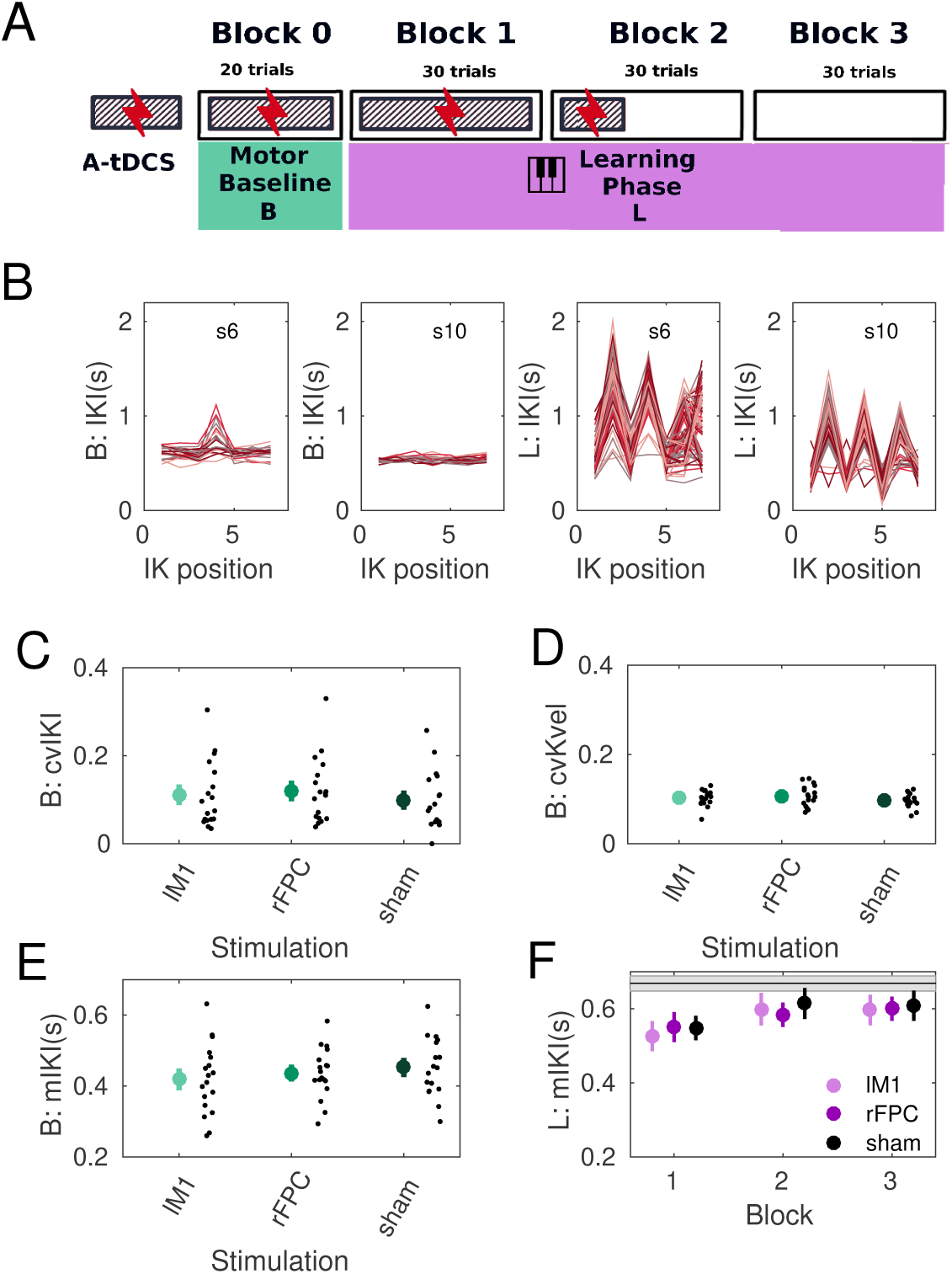
Experimental Design and Behavioral Results. **(A)** Participants were tested on three separate weeks during which either an active tDCS protocol over the lM1, or the rFPC, or a sham stimulation condition were applied. All tDCS protocols extended for 20 minutes, which included (i) an initial 3 minute resting phase, (ii) a baseline phase of regular isochronous motor performance, and (iii) part of the reward-based learning blocks (1 and 1/3). **(B)** Illustration of timing performance with sham stimulation during the baseline phase (B; panels 1-2) and learning phase (L; panels 3-4) in two participants. Timing was measured using the inter-keystroke-interval (IKI) in seconds, and shown for each inter-keystroke position (1 to 7 for sequences of 8 key presses). Different trajectories denote performance in different trials. **(C-E)** Effects of stimulation conditions on performance variables during the baseline phase (B). Colored big dots indicate means, with error bars denoting *pm* SEM. **(C)** Across-trials temporal variability, measured with the coefficient of variation of IKI or cvIKI; **(D)** across-trials variability in keystroke velocity or loudness, cvKvel; **(E)** mean performance tempo, mIKI (s). **(F)** Average performance tempo during learning (L). The gray horizontal line indicates the mean tempo of the hidden target solutions.

Next, during the reward-based learning blocks, participants had to play one sequence of eight or seven notes, which was a combination of the four neighboring white keys they had pressed during the baseline phase (**Figure S1**). Three different types of sequences were used for each stimulation session, with a pseudo-randomized counterbalanced order across participants. Each sequence was defined over a similar range of semitones but the range had a different spatial location on the keyboard (i.e. towards higher or lower pitch values, **Figure S1**). After completing the baseline phase, participants were explicitly taught the sequence key order by one of the experimenters, who played the melody using an isochronous timing. They were however instructed that the timing of the performance target was not isochronous and thus their goal was to use trial-based feedback (scores from 0 to 100) to approach the target timing of the performance.

The performance measure that was rewarded was the Euclidean norm of the vector corresponding to the pattern of temporal differences between adjacent inter-keystroke-intervals (IKI, in s) for a trial-specific performance (See Supplementary Materials). To approach the hidden target performance, participants had to deviate from an isochronous performance and find out the right combination of successive IKIs. Notably, however, different combinations of IKIs could lead to the same IKI differences and thus same Euclidean norm. Thus, the solution space was consistent with different movement choices obtaining the same reward. Accordingly, higher overall exploration would be associated with the perception that the environmental volatility was higher. This was explicitly assessed below in our mathematical model of the behavior.

The focality and magnitude of the neuromodulatory effects induced by the active tDCS protocols was assessed in each participant using electric field simulations (See *Materials and Methods*). The simulations were carried out with SimNIBS 2.1 software [30, 31] and using individualized head models obtained from the structural T1-weighted MR images. This analysis revealed that the focus the induced electric field was within the targeted regions and had a similar magnitude in both structures (Figure 2). The peak values of the vector norm of the electric field (normE) did not differ between active stimulation conditions (99.9% percentile: mean and SEM for lM1-tDCS = 0.132 [0.006] V/m; for rFPC-tDCS = 0.134 [0.010] V/m; permutation test, *P* > 0.05). In addition, the volume corresponding with the 99.9% percentile of the field strengh was not significantly different between active tDCS conditions (focality: 1.22 [0.11] × 10^4^mm^3^ for lM1-tDCS; 1.14 [0.08] × 10^4^mm^3^ for rFPC-tDCS; *P* > 0.05). Notwithstanding the similarity in peak and focality of the simulated normE values for lM1 and rFPC-tDCS, in both cases the electric field spread to neighboring areas beyond the target coordinate. Under lM1-tDCS, the induced electric field was maximum in lM1 (area 4 of the human connectome project multimodal parcellation, HCP-MMP1; [32]), followed by the premotor cortex (6), prefrontal areas (8Av and 8C) and somatosensory cortex (3). Under rFPC-tDCS, the peak of the electric field corresponded with the rFPC (areas 10p and 10pp), followed by regions in the medial prefrontal cortex (9) and orbitofrontal cortex (11). Lastly, the variability in the electric field strength (standard deviation) did not differ between tDCS targets (*P* > 0.05, **Figure S2**), supporting the comparable effects of both stimulation protocols in our sample.

**Figure 2:**
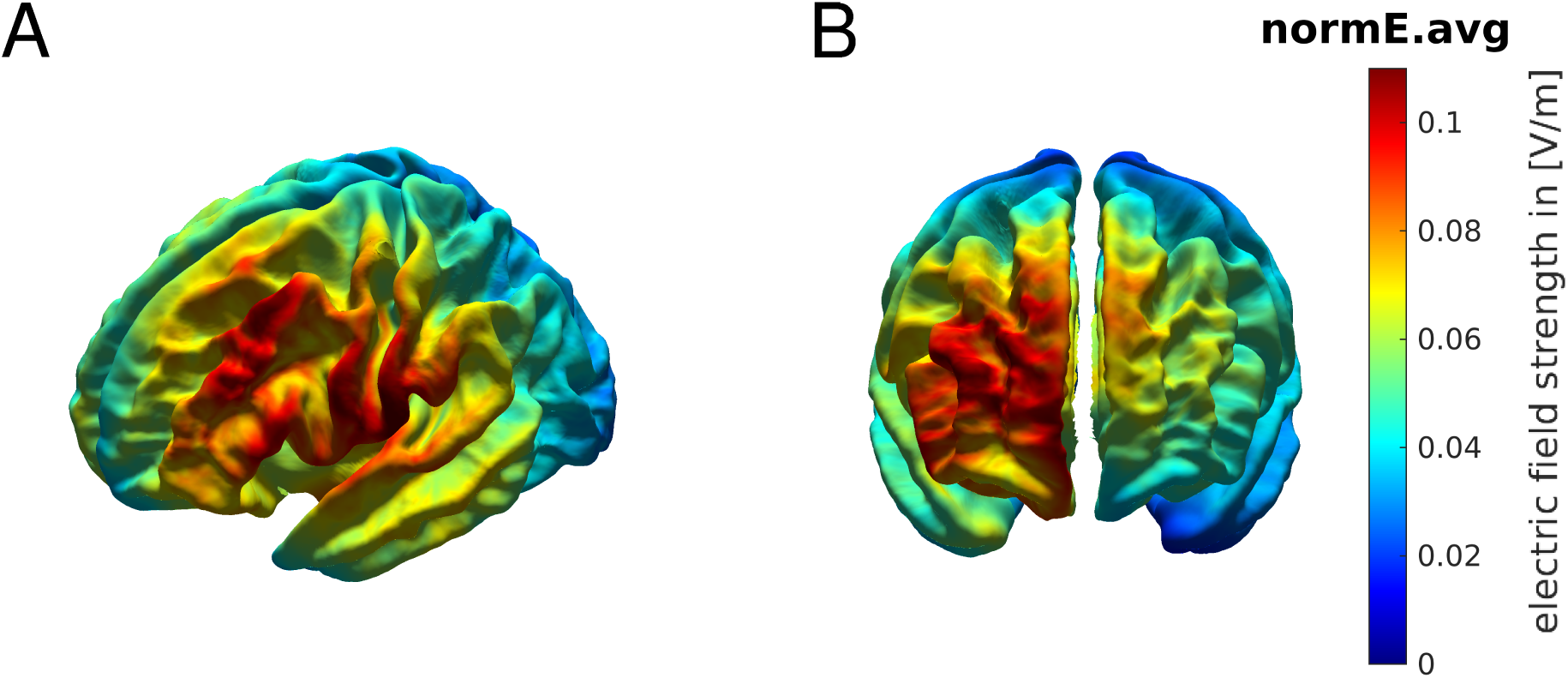
Electric field distribution for anodal lM1-tDCS (A) and rFPC-tDCS (B). Norm of the electric field strength (normE) derived from FEM calculations using SimNIBS, and averaged across participants using the fsaverage-transformed surface.

### Behavioral results

Analysis of the behavioral data focused on the assessment of motor variability across trials and along two different dimensions: time, which was the *instructed* task-related dimension and measured here using the IKI(s) index; and keystroke velocity (Kvel), which was the non-task-related dimension and is associated with the loudness of the key press. Motor variability in each dimension was measured using the coefficient of variation (cv = sd/mean). During the baseline phase cvIKI and cvKvel provided a measure of each participant’s motor noise. During learning blocks, the level of task-related motor variability, cvIKI, was considered to reflect both intentional exploration of this parameter and unintentional motor noise. Assuming an additive model of unintended (noise) and intended temporal variability contributing towards the cvIKI values expressed during learning blocks [4], we estimated the degree of task-related exploration simply by subtracting the cvIKI values of the baseline phase from the corresponding ones in the learning phase: cvIKI_explor_ = cvIKI_L_ − cvIKI_B_.

The achieved scores and other general performance variables, such as mean tempo (mIKI) and mean Kvel were also evaluated. During the baseline phase we assessed statistical differences between each pair of active and sham stimulation condition in the abovementioned variables, excluding the scores and cvIKI_explor_. This was done with pair-wise permutation tests for matched samples [33]. Further, during the learning blocks, statistical analysis of all dependent variables was performed separately for each active stimulation condition relative to sham using a 2 × 3 non-parametric factorial analysis with factors Stimulation (rFPC, sham; or lM1, sham) and learning Block (1,2,3). This analysis was implemented using synchronized rearrangements [34]. As additional planned analysis, we evaluated the change from block 1 to 3 in all dependent variables and contrasted them between active and passive stimulation conditions using permutation tests.

The different active tDCS protocols did not have dissociable effects on baseline motor variability for timing when compared to sham (cvIKI; *P* > 0.05 in both cases; Figure 1). Neither was there a significant difference in cvKvel at baseline between active and sham stimulation conditions (*P* > 0.05). General performance parameters, such as the mean tempo or keystroke velocity, did not differ in this phase as a function of the stimulation protocol either (*P* > 0.05, for mIKI and mKvel).

During reward-based learning participants improved their scores across blocks in all tDCS conditions (Figure 3A; main effect Block, P = 0.0001, using rFPC and sham for factor Stimulation; similar result for lM1 and sham: main effect Block, P = 0.002). These data support that participants successfully used the trial-wise feedback in all tDCS conditions to learn about the hidden goal. There was no main effect for Stimulation or Interaction effect in either factorial analysis (*P* > 0.05). Planned pairwise comparisons between the change in scores from block 1 to 3 for each active tDCS condition and sham demonstrated a significantly larger increase following rFPC-tDCS relative to sham (Figure 3B; increase of 19.9 [standard error of the mean or SEM 3.77] for rFPC-tDCS; increase of 12.3 [2.40] for sham: P = 0.047; moderate effect size, assessed with a non-parametric effect size estimator for dependent samples [35], Δ_*dep*_ = 0.62, confidence interval or CI = [0.50, 0.80]); no differences between lM1 and sham were found (*P* > 0.05; increase of 12.6 [3.8] for lM1-tDCS).

**Figure 3:**
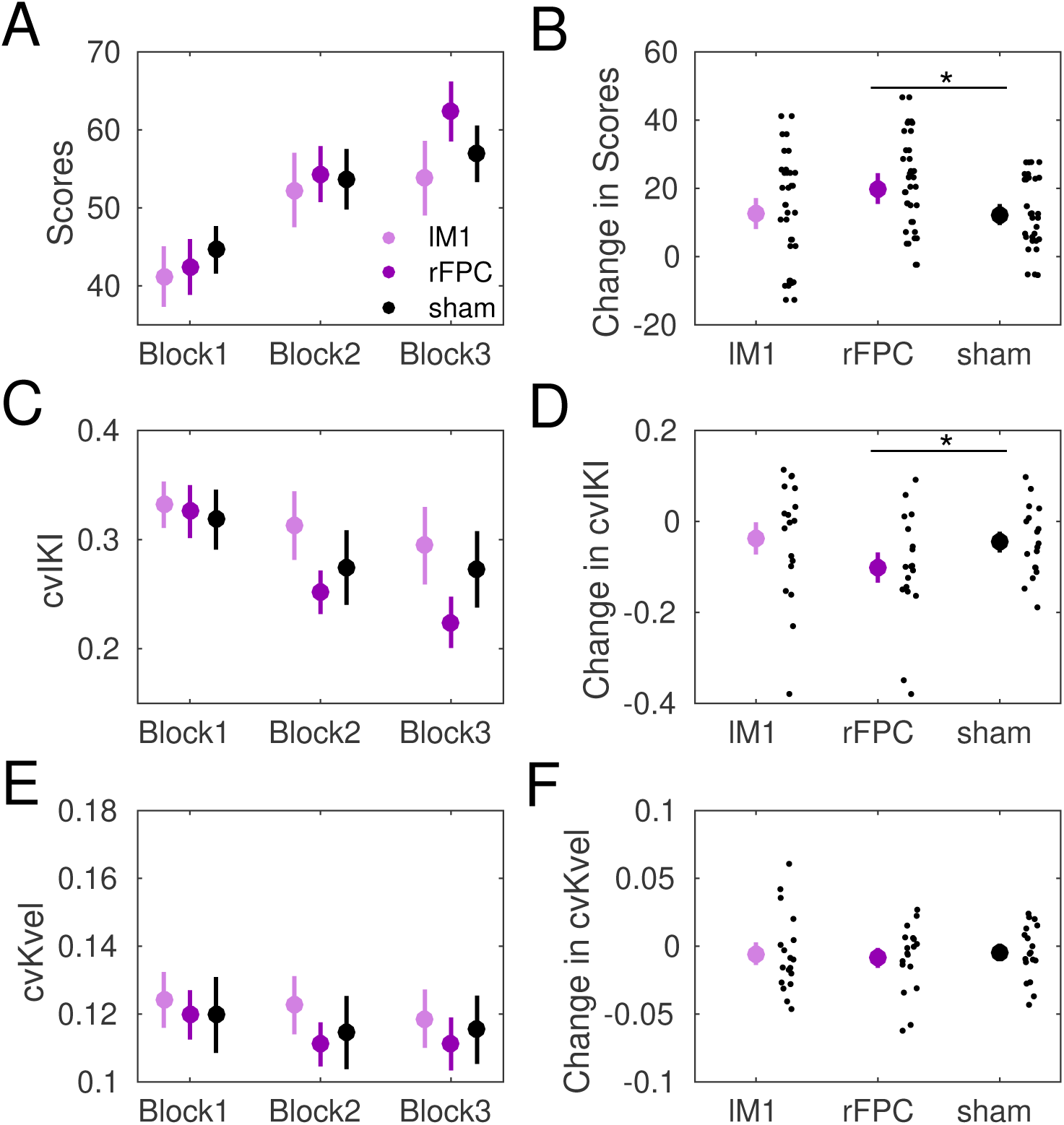
Behavioral Results During Reward-Based Learning. **(A)** Participants increased their scores across blocks regardless of the stimulation condition (significant main effect Block in both separate factorial analyses using factors Block (1-3) and Stimulation (rFPC, sham: or lM1, sham), supporting they successfully used the trial-by-trial feedback to approach the hidden performance goal. **(B)** The change in scores from block 1 to 3 was significantly larger with rFPC-tDCS than sham (denoted by the asterisk; P = 0.047, permutation tests; moderate effect size Δ_*dep*_ = 0.62, CI = [0.50, 0.80]). **(C)** Same as (A) but for the degree of temporal variability, cvIKI, see main text for results. **(D)** The reduction from block 1 to 3 in cvIKI was significantly more pronounced in rFPC-tDCS than sham (P = 0.041, Δ_*dep*_ = 0.68, CI = [0.50, 0.83]). No significnat effects were found when comparing lM1-tDCS and sham. **(E-F)** Same as (C-D) but for keystroke velocity. No significant main effects or interactions were found when assessing cvKvel. Neither were there differential effects of stimulation on the change in cvKvel from block 1 to 3. Small black does represent individual participant data. Colored dots display mean values with error bars denoting ± SEM.

The general increase in scores across blocks was paralelled by a reduction in the expression of task-related motor variability, measured using cvIKI (Figure 3C; significant main effect of Block in both factorial analyses: P = 0.0069 for rFPC and sham; P = 0.023 for lM1 and sham). No significant main effect for Stimulation or interaction effect was found. Planned pairwise analyses revealed that following rFPC-tDCS the drop in temporal variability from block 1 to 3 was more pronounced than following sham (P = 0.041, Δ_*dep*_ = 0.68, CI = [0.50, 0.83]). When comparing lM1-tDCS to sham, the reduction in motor variability was not significantly different (*P* > 0.05). Next, we assessed the effect of tDCS on the index of task-related motor exploration cvIKI_explor_. This measure also displayed a significantly larger reduction from block 1 to 3 following active rFPC stimulation compared to sham (P = 0.024, Δ_*dep*_ = 0.75, CI = [0.50, 0.94], larger effect size). This outcome supported that rFPC-tDCS relative to sham contributed to the increase in scores across blocks primarily through a regulation of exploration, without concurrent modulation of motor noise (Figure 4).

**Figure 4:**
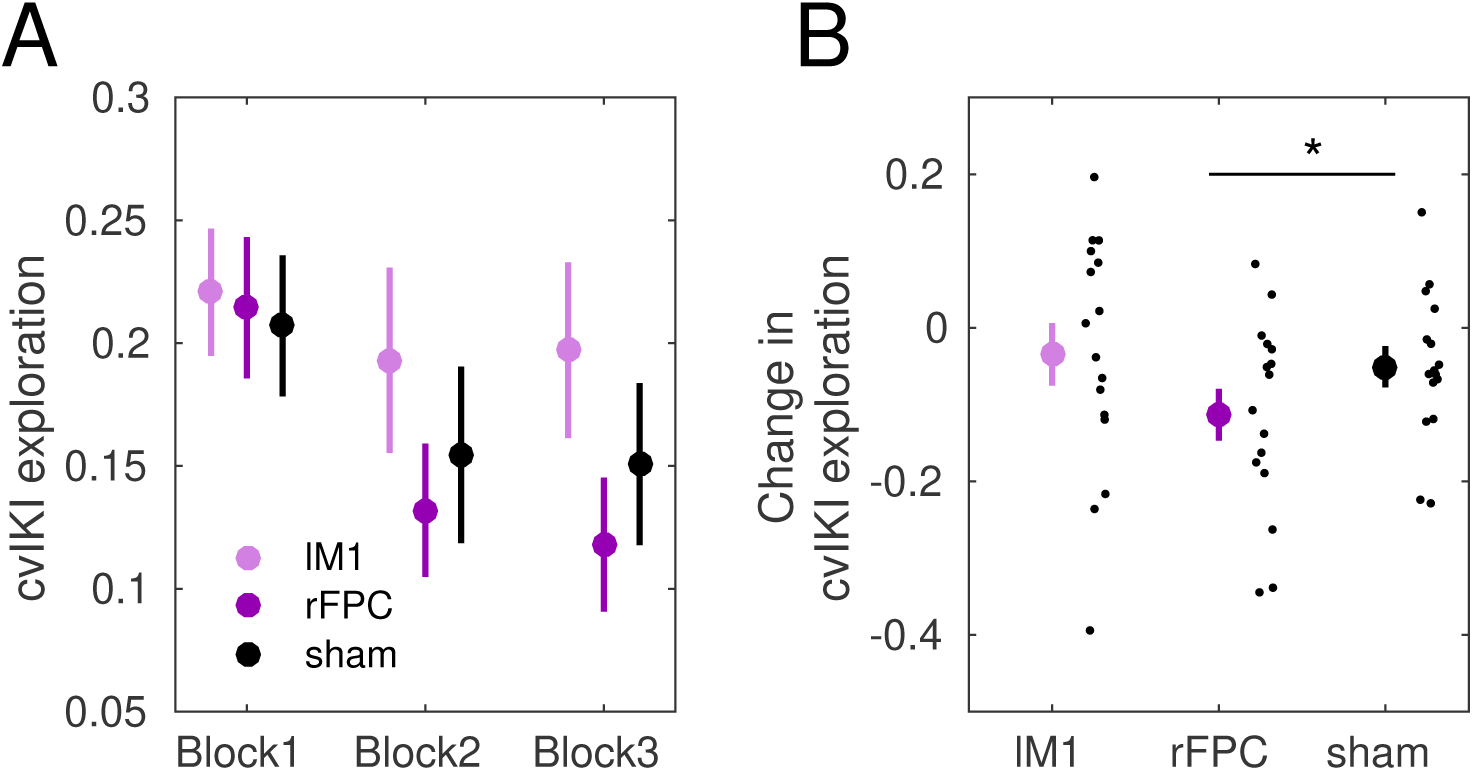
Exploration during Reward-Based Learning. **(A)**. We assumed an additive model of unintended (noise) and intended temporal variability contributing towards the cvIKI values obtained in learning blocks. Accordingly, to obtain the component of variability related to task-related exploration during learning, cvIKI_explor_, we simply subtracted the cvIKI index of the baseline and learning phases: cvIKI_L_ − cvIKI_B_. **(B)** There was a significantly larger drop from block 1 to 3 in cvIKI_explor_ following active rFPC stimulation compared to sham (P = 0.024, permutation tests; Δ_*dep*_ = 0.75, CI = [0.50, 0.94], large effect size). Small black dots show individual participant values; colored big dots represent means, with error bars denoting ± SEM.

Finally, control analyses carried out on non-task-related variables, revealed no significant main effects or interactions on the variability in keystroke velocity (Figure 3E), yet a main effect of factor Block on the mean performance tempo (mIKI; P = 0.036 for Stimulation rFPC-sham; P = 0.042 for Stimulation lM1-sham). This effect reflected the observed increase in performance tempo across blocks (Figure 1F), which was related to the overall slower timing of the rewarded solution relative to their initial assumption. The change in mean tempo from block 1 to 3, however, did not differ between active and sham stimulation conditions (*P* > 0.05). On average, participants played the sequences at a rate of one keystroke every 0.57 (0.036) s under lM1-tDCS, 0.58 (0.027) s under rFPC-tDCS, and 0.59 (0.031) s during sham.

### Bayesian model of learning

We investigated how individuals used the reward feedback to infer the hidden performance goal using a hierarchical Bayesian model, the Hierarchical Gaussian Filter (HGF, [36, 27]). The HGF was adapted to model participants’ beliefs about the reward on the current trial *k*, 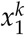, and about its rate of change, termed environmental volatility 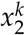. Volatility emerged from the multiplicity of performance-to-score mappings, as different temporal patterns of the performance with identical IKI-difference values led to the same scores. Beliefs on *x*_1_ and *x*_2_ were Gaussian distributions and thus fully determined by the sufficient statistics *µ*_*i*_ (*i* = 1, 2, mean of the posterior distribution for *x*_*i*_, corresponding with participants’ expectation) and *σ*_*i*_ (variance of the distribution, representing uncertainty of the estimate). The belief trajectories about the external states *x*_1_ and *x*_2_ (mean, variance) were further used to estimate the most likely response corresponding with those beliefs. A schematic illustrating the model structure and outputs is shown in Figure 5.

**Figure 5:**
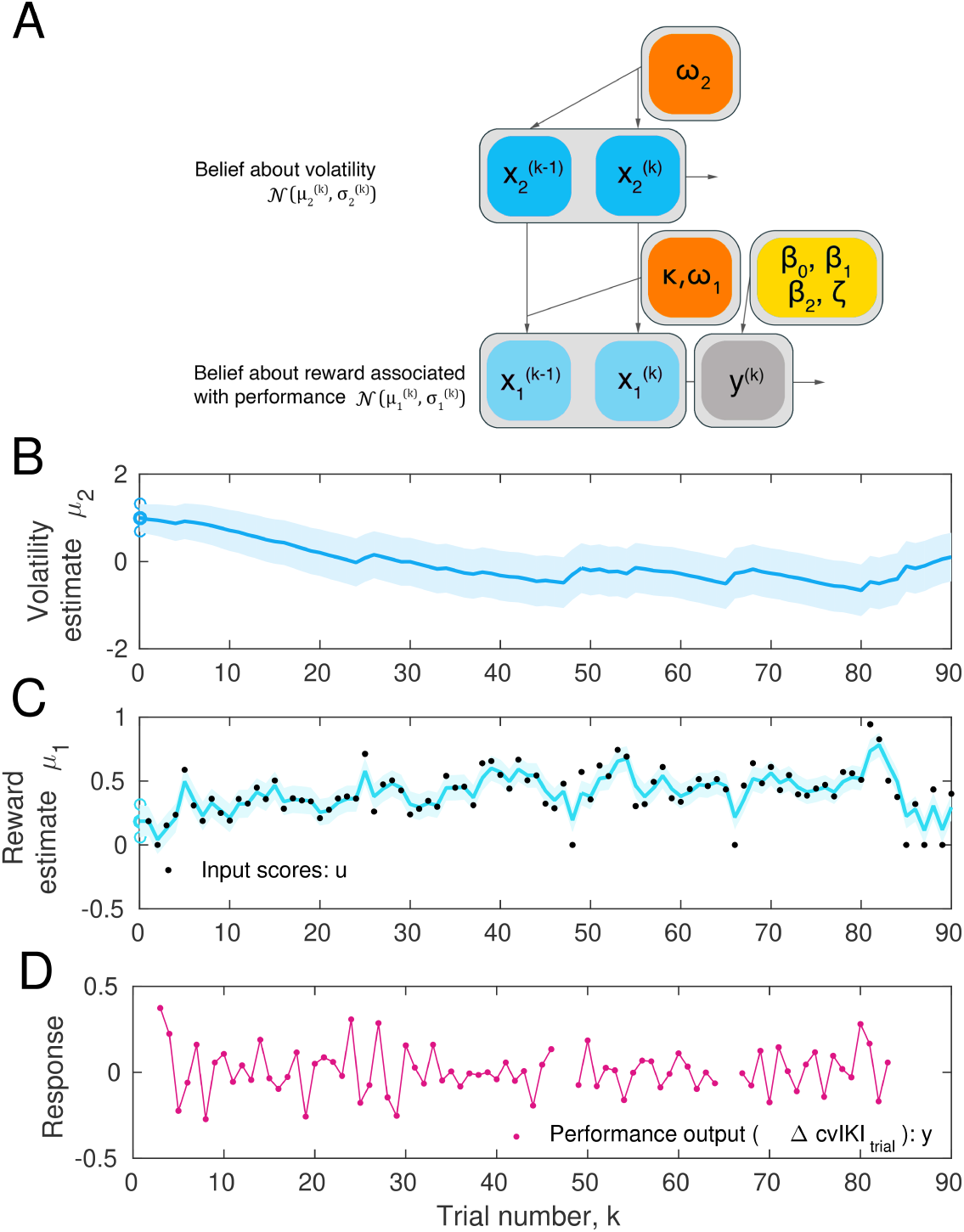
Computational Model. Two-level Hierarchical Gaussian Filter for continuous inputs. **(A)** Schematic of the two-level HGF, which models how an agent infers a hidden state in the environment (a random variable), *x*_1_, as well as its rate of change over time (*x*_2_, environmental volatility). Beliefs (*µ*_1_, *µ*_2_) about those hierarchically-related hidden states (*x*_1_, *x*_2_) at trial k are updated with the input scores for that trial via prediction errors (PEs). The states *x*_1_ and *x*_2_ are continuous variables evolving as coupled Gaussian random walks, where the step size (variance) of the random walk depends on a set of parameters (shown in orange boxes). The lowest level is coupled to the level above through the variance. The response model generates the most probable response, *y*, according to the current beliefs about the lower state, *x*_1_, and is modulated by the response model parameters (yellow box).**(B)** Example of trial-by-trial beliefs about volatility, *µ*_2_. **(C)** Belief on the first level, which represents an individual’s expectation of reward in trial *k, µ*_1_. Black dots represent the trial-wise input scores (*u*). **(D)** Performance output (change in trial-wise cvIKI_trial_; see main text). Shaded areas denote the variance or informational uncertainty on that level.

The response model defines the mapping from the trajectories of perceptual beliefs onto the observed responses in each participant. We were interested in assessing how belief trajectories or related computational quantities influenced subsequent behavioral changes, potentially increasing trial to trial exploration or rather driving exploitation. Accordingly, we constructed different response models corresponding with different scenarios in which participants would link a specific performance measure to reward, such as the sequence duration or the degree of temporal variability (See *Materials and Methods*).

Each model was fitted with the 90 trial-wise performance values of the response model and the input scores for each tDCS session, and the log model-evidence was used to optimize the model fit [37]. Random Effects Bayesian Model Selection (BMS; [38]; code freely available from the MACS toolbox, [39]) selected as best model the one that explained the change in the trial-wise coefficient of variation of adjacent IKI values (cv across sequence positions in one trial; termed cvIKI_trial_ to dissociate it from the measure of across-trials variability, cvIKI): 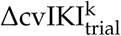. This variable reflected changes between previous *k* − 1 and current *k* trial in cvIKI_trial_, with greater changes indicating more exploratory behavior along this parameter from trial to trial. In the winning model, this performance variable was explained as a linear function of precision-weighted prediction errors (pwPE) about reward, termed *ϵ*_1_, and volatility, termed *ϵ*_2_, on the preceeding trial *k* − 1:

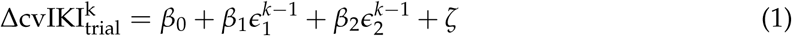

where *ζ* is a Gaussian noise variable. The rationale for choosing pwPE at level 1 and 2 as predictors in some of the alternative response models was the relevance of PE weighted by uncertainty in current frameworks of Bayesian inference. Moreover, pwPEs determine the step size of the update in the expectation of beliefs: 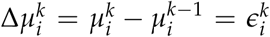. That is, larger pwPE about reward increase the expectation of reward, while larger pwPE about volatility increase the corresponding volatility estimate.

All *β* coefficients of the multiple linear regression response model were significantly different than zero in each stimulation condition (*P*_*FDR*_ < 0.05), with the exception of *β*_0_ for rFPC-tDCs and *β*_2_ for sham. *β*_0_ was positive and comparable in lM1 and sham stimulation conditions (*P* > 0.05, and it was significantly larger in sham relative to rFPC-tDCS (*P*_*FDR*_ < 0.05). Further, on average *β*_1_ was negative in each active and sham stimulation condition. This outcome indicated that larger pwPE about reward in the previous trial (contributing to a higher expectation of reward) promoted an attenuation in the change of the performance variable 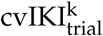. Moreover, rFPC-tDCS decreased the sensitivity of this association (reduced “negative” slope, larger *β*_1_ values in rFPC than sham, *P*_*FDR*_ < 0.05). The effect of pwPE about volatility on trial to trial behavioral changes was also dissociated between rFPC-tDCS and sham stimulation, as *β*_2_ coefficients were negative and significantly smaller for rFPC relative to sham stimulation (*P*_*FDR*_ < 0.05). Accordingly, larger pwPE about volatility in the previous trial—related to an increase in the expectation of volatility—contributed to subsequent exploitation for rFPC-tDCS. By contrast, during sham stimulation this association was not significant (*P* > 0.05), suggesting that pwPE about volatility did not consistently modulate trial-to-trial behavioral changes. Lastly, the increase in 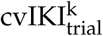 following larger pwPE on volatility on the previous trial was enhanced for lM1-tDCS relative to sham (larger positive *β*_2_ during lM1-tDCS, *P*_*FDR*_ < 0.05). Thus, the winning response model identified different behavioral strategies in response to updates in reward and volatility as a function of the tDCS stimulation condition.

The HGF with the winning response model provided a good fit to the behavioral data, as the examination of the residuals shows(**Figure S3**). There were no systematic differences in the model fits across tDCS conditions. Additionally, estimation of belief trajectories using different types of simulated responses revealed that agents observing a broader range of outcomes or introducing more trial to trial behavioral changes have higher expectation on volatility, as both contributed towards an increased rate of change in reward (= volatility; **Figure S4**). Additional details are provided in *Supplementary Materials*.

Using the winning model, we further examined the effects of Stimulation ([rFPC, sham] and separately [lM1, sham]) and Block (1-3) on the Bayesian learning process by submitting the relevant model parameters to two separate 2 x 3 nonparametric factorial analyses. The main variables for the statistical analysis were the mean trajectories of perceptual beliefs (*µ*_1_, *µ*_2_), and the estimation uncertainty about those beliefs (variances *σ*_1_, *σ*_2_; note that the inverse variance is termed precision, corresponding with the confidence placed on those beliefs). The factorial analyses were followed up by between-stimulation pairwise permutation tests, similarly to the model-free analysis.

For rFPC and sham, we found a significant main effect of factor Block on the belief estimates about reward (*µ*_1_, P = 0.002), volatility (*µ*_2_, P = 0.0012; Figure 6) and the uncertainty about volatility (*σ*_2_, P = 0.0009; **Figure S5**). Across blocks, the expectation of reward increased, whereas the volatility estimate decreased, as participants gradually shifted to exploitation and approached the hidden performance goal (Figure 3). In addition, there was an interaction effect for *µ*_1_ (P = 0.0002), as this estimate increased more steeply with rFPC-tDCS than sham. Planned statistical analyses of differences between conditions separately at each block demonstrated that the expectation of reward was higher during the last block for rFPC-tDCS relative to sham (*P* < *P*_*FDR*_; effect size Δ_*dep*_ =0.70, CI = [0.56, 0.83]). Moreover, volatility estimates were smaller following rFPC as compared to sham during the initial learning block (*P* < *P*_*FDR*_; Δ_*dep*_ = 0.68, CI = [0.56, 0.80]). Combining these findings with the result of negative *β*_1_ coefficients in the response model (Figure 6D), it follows that the increased expectation of reward in rFPC-tDCS driven by large pwPE values on this level was associated with a more pronounced tendency to exploit the inferred solution (smaller change in cvIKI_trial_).

**Figure 6:**
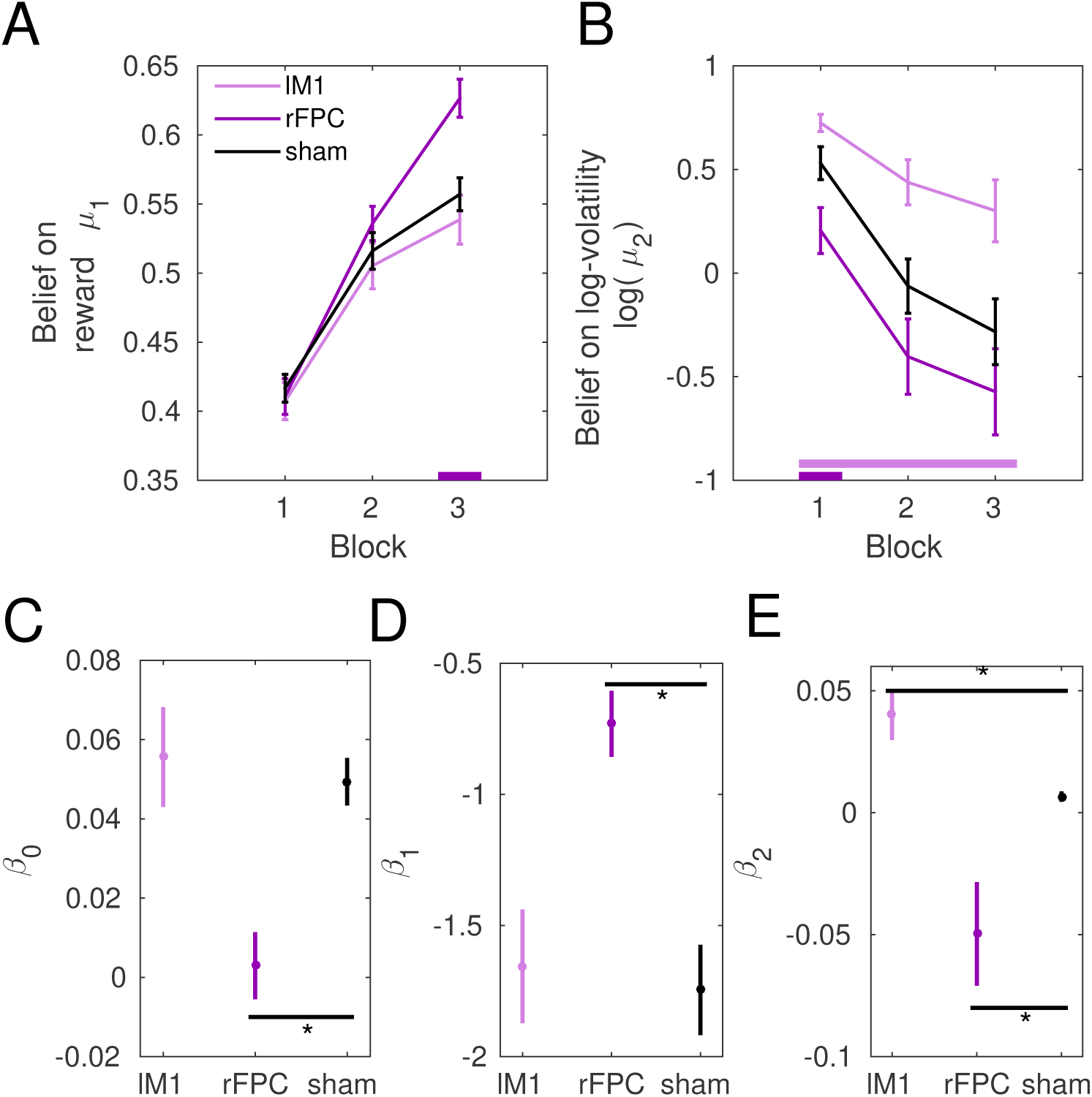
Computational modeling analysis. Data shown as mean and ± SEM. **(A)**. Belief trajectories for reward. Participants under rFPC-tDCS had a higher expectation on reward, *µ*_1_, corresponding with their higher achieved scores (*P* < *P*_*FDR*_; effect size Δ_*dep*_ =0.70, CI = [0.56, 0.83], dark purple bar at the bottom). **(B)** Trajectories of beliefs on volatility. The expectation on environmental (phasic) log-volatility (log(*µ*_2_)) was significantly larger in the initial block following rFPC-tDCS relative to sham stimulation (*P* < *P*_*FDR*_; Δ_*dep*_ = 0.68, CI = [0.56, 0.80]). This quantity was larger during lM1-tDCS relative to sham (*P* < *P*_*FDR*_; Δ_*dep*_ = 0.71, CI = [0.58, 0.82]).**(C-E)**. *β* coefficients of the winning response model that explain the performance measure in trial *k* as a linear function of a constant value (intercept: 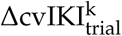), and the belief estimates for reward and log-volatility on the previous trial: 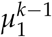 and 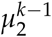. The performance measure in the winning model, 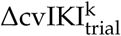, was the change in the degree of temporal variability across keystroke positions from trial *k* − 1 to *k*. The *β* values of the winning response model are plotted separately in each stimulation condition. Differences between active and sham stimulation conditions in *β* coefficients are denoted by the horizontal black line and the asterisk (*P*_*FDR*_ < 0.05, see main text).

For lM1 and sham, there was a significant main effect of factor Block on the belief estimates about reward, volatility, as well as on the uncertainty about volatility (P = 0.005, 0.001 and 0.005; respectively). In addition, there was a significant main effect of factor Stimulation for volatiluty estimates, *µ*_2_ (Figure 6B), and their uncertainty *σ*_2_ (**Figure S5**). Planned analyses using pair-wise permutation tests between conditions for each separate block further revealed that the expectation of volatility was higher under lM1-tDCS than sham (*P* < *P*_*FDR*_; Δ_*dep*_ = 0.71, CI = [0.58, 0.82]). Participants under lM1 stimulation therefore believed the reward tendency across trials was more unstable. They also were more uncertain about their knowledge of reward (higher uncertainty estimate *σ*_1_ in blocks 1-3, *P* < *P*_*FDR*_; Δ_*dep*_ = 0.67, CI = [0.55, 0.79]). Given the higher positive *β*_2_ coefficients in the response model obtained for lM1 relative to sham stimulation, these results indicate that larger pwPE about volatility (leading to increased *µ*_2_ in lM1-tDCS) contributed more to increasing the measure of temporal variability in the next trial.

## Discussion

In this study, we identify a crucial role of rFPC in reward-based motor learning by using a motor task adapted from Bayesian learning settings [36, 27], a Bayesian computational model of the behavior, and simulations of the electric field induced by each tDCS target. The results indicated that rFPC-tDCS relative to sham improved the scores across learning blocks primarily through a regulation of task-related exploration, without concurrent modulation of motor noise. In contrast to the salient effect of rFPC-tDCS on reward-based motor learning, lM1-tDCS modulated behavior in a manner that was not significantly different from sham stimulation. The outcomes extend findings in the area of decision-making [13, 12, 11, 10] to that of motor skill learning: Brain regions previously linked to decision-making in the cognitive or perceptual domain, such as the FPC, could be relevant in the motor domain.

Further, both active tDCS protocols led to an increase in the simulated electric field strength, corresponding with enhanced local plasticity as shown in a recent study assessing the neurophysiological effects of tDCS with SimNIBS and magnetic resonance spectroscopy [40]. Accordingly, a mechanism by which rFPC-tDCS may regulate exploration comes about through an enhanced recruitment of the local neuronal populations in the rFPC region. An important implication of the findings is that the rFPC might be a good target for noninvasive brain stimulation in motor learning and rehabilitation settings that use reinforcement or reward signals.

### Role of frontopolar cortex in the intentional modulation of motor variability

Neuromodulation of the rFPC via anodal tDCS reduced the use of motor variability during learning blocks. This effect was accounted for by a decrease in the intentional use of motor variability —exploration —after controlling for changes in motor noise. Recent work has demonstrated that the inhibition of the rFPC via TMS during binary choices decreased exploration, but not random exploration [10]. The rFPC has also been shown to be central for regulating the balance between exploration and exploitation in a three-armed bandit task [11]. Our results expand these findings to the area of motor learning, demonstrating that anodal rFPC-tDCS regulated the intentional use of motor variability by promoting exploitation of the movement inferred to be highly rewarding.

The outcome of anodal rFPC-tDCS promoting exploitation seems to stand in contrast with the anodal rFPC-tDCS effects reported in [11], leading to increased exploration of alternative bandits. Evidence from our Bayesian computational model helps clarify this aspect of the results. We found a pronounced drop in the volatility estimate under rFPC-tDCS, which meant that participants perceived the reward tendency to be more stable than during sham. In the continuous movement space over which our task was defined, timing was the task-related dimension that needed to be mapped to reward. Thus, inferring this complex mapping might have been facilitated by rFPC-tDCS. This is supported by the reduced volatility estimates which showed that participants had an expectation of a more stable (reduced change in) reward tendency across trials. In addition, rFPC-tDCS made participant’s trial-to-trial behavioral changes more sensitive to update steps in volatility, an effect that was not observed during sham stimulation. More specifically, larger pwPE about volatility contributed towards exploitation of the current behavioral response on the following trial. By contrast, during lM1-tDCS large pwPE about volatility, and therefore an increase in volatility estimates, promoted further exploration. The results are consistent with previous findings indicating that FPC managed competing goals by keeping track of alternative choices [13, 12]. Moreover, FPC would also infer the absolute reliability of several alternative choices [41, 42], which is consistent with the Bayesian results on reduced volatility. Notably, our results expand previous findings by revealing that during motor learning, which is generally solved in a continuous movement space [6], learning the mapping between different movement configurations and their associated reward relies on FPC, given its contribution to facilitating assessment of the relative value of multiple alternative choices [41]. In addition, the findings support that lM1 and rFPC-tDCS have opposing effects on how behavioral adaptations during learning rely on volatility estimates. While trials increasing the expectation of volatility induced subsequent exploration under lM1-tDCS, they led to exploitation of the current motor programme under rFPC-tDCs.

Stimulation over the contralateral M1 did not modulate reward-based motor learning in a significantly different way than the sham stimulation when using model-free analysis. This lack of significant effects should be interpreted with caution as our statistical approach does not allow us to make inferences on null results. The outcome was, however, not entirely unexpected, as standard M1-tDCS protocols generally assess effects on motor retention or consolidation [43, 19, 44, 15] and have so far not reported modulations of motor learning processes that depend on reward signals. One notable exception is a recent anodal tDCS study that used smaller electrodes anterior and posterior to M1 [28]. This work demonstrated a benefit of the combination of reward signals and M1-tDCS on motor retention during a motor adaptation paradigm. This study did not, however, identify a faster learning rate under M1-tDCS, which was the effect we observed in the rFPC-tDCS protocol. The authors in [28] argued that reward signals combined with tDCS may influence motor retention through long-term potentiation mechanisms. It is therefore likely that an effect of M1-tDCS on reward-based motor learning primarily affects retention of motor skills and not initial learning, as our task was designed to assess. More consistent evidence for a role of M1 in reward processing comes from TMS paradigms. In particular, M1-TMS studies of value-based decisions have shown an increase in motor cortex excitability that scaled with the anticipated reward value of a choice [23, 24]. Additional TMS work demonstrated an increase in motor cortex excitability during motor preparation in anticipation of reward delivery, thus providing compelling evidence for an involvement of M1 in reward-guided motor processing [21, 22]. This effect could be mediated by ventral striatal reward circuitry, given its established role in instrumental learning for reward [45, 46]. Because of the null results of M1-tDCS in our task, resolving the issue of the dissociable role of M1 and FPC in regulating motor variability during reward-based motor learning will require follow up TMS studies and a comparison between initial learning and motor retention [47].

Of note, a limitation of any tDCS study is that the diffuse spatial effect of tDCS does not allow us to determine whether modulation of the target area alone is responsible for the observed behavioral effect [48]. Even with the higher spatial resolution of TMS, it has been argued that any effect of FPC stimulation on behavior could be mediated by other brain regions coupled with FPC during behavioral exploration [10], such as the inferior parietal cortex or ventral premotor cortex [41] or by regions engaged in other aspects of goal-directed behavior, such as the ventromedial prefrontal cortex and orbitofrontal cortex [25, 12].

To mitigate that limitation, we carried out electric field simulations in the individual anatomy and assessed the strength and focality of the induced electric field for each tDCS target. The simulations demonstrated an enhanced focal activation in the targeted area, corresponding with increases in activation (excitatory effects [40]) and with a focality and peak value not significantly different between both active stimulation protocols. However, the induced electric field also extended to neighboring regions, such as the OFC and the medial PFC for rFPC-tDCS and premotor and somatosensory areas for lM1-tDCS. The latter resemble recent findings in lM1-tDCS with SimNIBS [40]. Notwithstanding the certain degree of diffusion of the induced electric field, we could establish that the spatial effects of tDCS for both protocols were consistent with the target area. Future studies combining electroencephalography and functional MRI should assess the network of interactions between FPC, other regions in the prefrontal cortex, and cortical motor regions to determine the precise mechanism underlying the rFPC-tDCS effect on reward-based motor learning reported here.

## Materials and Methods

### Participants

Nineteen right-handed participants (10 females, mean = 27.7 yrs, std = 3.3 yrs, range 21-33) with no musical training were recruited. Laterality quotient was assessed by the Oldfield handedness inventory ([49]; mean = 90, SEM = 3.2; values available in 17/19 participants). The sample size is similar to that found in tDCS studies focusing on motor learning [50, 28]. Participants had no history of neurological disease or hearing impairment. They gave written informed consent before the beginning of the first experimental session. The study protocols were approved by the local ethics committee of the of the University of Leipzig (277-14-25082014). To incentivize participants during completion of the reward-based learning phase of the motor task, they were informed about a Â£50 voucher for online purchases that would be awarded to the participant scoring the highest average points across the three sessions.

### Experimental Design

A sham-controlled, double-blinded, cross-over design was implemented. The study was comprised of three sessions with a 7-days interval between them (same time of day) to reduce potential carry-over effects. Study procedure for each session was identical, with active or sham tDCS applied to the target area for a period of 20 min during task performance (Figure1). Participants received either right FPC or left M1 anodal tDCS, or sham tDCS (targetting either rFPC or lM1, 50% − 50% split across participants). Sham, rFPC and lM1 stimulation conditions were counterbalanced across participants. More details are provided in the *tDCS* section.

### tDCS

During the experiment, tDCS was applied via saline-soaked sponge electrodes to the individual target coordinate using a battery-driven DC-stimulator (NeurConn, Ilmenau, Germany). tDCS can transiently modulate cortical excitability via application of direct currents, as shown in combined TMS-tDCS studies [18, 51], and further supported by recent simulation studies [40]. Anodal tDCS has been shown to increase cortical excitability, however additional evidence indicates that it could lead to the opposite polarity of the effect, reducing cortical excitability [48]. For instance, extending the stimulation duration beyond 26 min has been shown to result in inhibitory rather than excitatory effects after anodal tDCS [52]. To control for this confound, we complemented the main analysis with a simulation of the electric field induced by tDCS in each participant using SimNIBS (see below). Anodal or sham tDCS was applied to the right FPC or left (contralateral) M1 regions. The target coordinate for rFPC-tDCS was selected from previous tDCS and fMRI work investigating the role of rFPC on exploration (Montreal Neurological Institute or MNI peak: x = 27, y = 57, z = 6; [25, 11]). For lM1-tDCS, we used a target coordinate in the hand area of the left primary motor cortex (MNI peak: x = -37, y = -21, z = 58), based on [53]. The target coordinates were transformed to the individual native space using a T1-weighted high-resolution magnetic resonance image from each participant. Specifically, the MNI coordinates were converted into participants’s native MNI space using the reverse native-to-MNI transformation from Statistical Parametric Mapping (SMP, version SPM12). The point on the scalp corresponding with each of our targeted brain areas was marked to place the active electrode (5×5 cm^2^). The reference (cathode, 10 × 10 cm^2^) electrode for rFPC-tDCS was placed at the vertex [11], whereas it was located over the frontal orbit for lM1-tDCS [18]. Flexible elastic straps were used to fixate the electrodes on the head. A three-dimensional (3D) neuronavigation device (Brainsight Version 2; Rogue Research, Montreal, Canada) was used to guide positioning of active electrodes.

We stimulated with a weak direct current of 1 mA in all conditions for 20 minutes resulting in a current density of 0.04 mA/cm2 under the target electrode and 0.01 mA/cm2 under the reference electrode. In general, the modulatory effect of tDCS on brain excitability is more pronounced after several minutes and can subsequently outlast the stimulation duration for up to 1.5 h [18, 51]. To account for the potential delay in the effect of tDCS, we instructed participants to wait for 3 minutes before we initiated the motor task. At the start of the active tDCS stimulation the current was ramped up for 30 s to minimize the tingling sensation on the scalp, which generally fades over seconds [15], and was also ramped down for 30 s at the end. During sham tDCS, the current was ramped-up for 30 s, held constant at 1 mA for 30 s and ramped-down for 30 s. This procedure aimed to induce a similar initial tingling sensation in active and sham protocols, yet with no modulations of cortical excitability for sham tDCS [54]. Before and after each tDCS session, participants rated on a 1-10 scale their fatigue, attention and discomfort levels, as well as the sensation with tDCS (post-tDCS, scale 1-5). No significant differences between active and sham tDCS sessions were found in the reported levels of fatigue, discomfort or attention levels (*P* > 0.05, post minus pre changes). The sensation was not different between rFPC and sham tDCS, either (*P* > 0.05). However, lM1-tDCS induced a higher sensation than sham (P = 0.0034, paired permutation test, Δ_*dep*_ = 0.87, CI = [0.57, 0.88]).

### Acquisition of behavioral data

Performance information was saved as MIDI (Musical Instrument Digital Interface) data, which provided the time onsets of keystrokes relative to the previous event (inter-keystroke interval, IKI, s), MIDI note number that corresponds with the pitch, and MIDI velocity (related to loudness).

Behavioral data are available in the Open Science Framework Data Repository: https://osf.io/zuab8/

### Bayesian model of behavior

The HGF has been extensively used to model reward-based learning in volatile environments [55, 37]. It was introduced as a perceptual model to explain how an agent infers a hidden state in the environment (a random variable), *x*_1_, as well as its rate of change over time (*x*_2_, environmental volatility). The hidden states (*x*_1_, *x*_2_) are continuous variables evolving as Gaussian random walks coupled through their variance. Thus, their value at trial *k* will be normally distributed around their previous value at trial *k* − 1. The posterior distribution of beliefs about these states (*x*_*i*_ = 1, 2) is therefore fully determined by the sufficient statistics *µ*_*i*_ (mean) and *σ*_*i*_ (variance). The perceptual model can be further coupled with a response model to generate responses based on those inferred states. In the HGF, beliefs are updated given new sensory input or observed outcomes via prediction errors (PEs). PEs are weighted by the ratio between the uncertainty of the current level and the lower level. Larger uncertainty reflects that an individual knows less about a quantity she is trying to infer, thereby promoting greater updates in the corresponding belief estimate. We used the series of feedback scores as input to the HGF.

Here, we implemented eight versions of the HGF with alternative response models that explained different performance measures as a function of relevant HGF quantities. The eight models fall into two families of related models, with each family being associated with a different performance measure: (i) The trialwise coefficient of variation of successive IKI values (cv across sequence positions; termed cvIKI_trial_; N = 4 models); (ii) the trialwise logarithm of the mean performance tempo (log(mIKI_trial_), in ms; N = 4 models). Note that cvIKI_trial_ was associated with the reward function, as larger differences between adjacent IKI values led to higher reward and also largercvIKI_trial_. This variable was therefore a strong candidate for the response model. Additionally, we considered the logarithm of the mean performance tempo in the trial to assess whether participants speeded or slowed down in each trial without changing the differences between IKI patterns. We were particularly interested in relating HGF computational quantities in the previous trial *k* − 1 to changes in performance from trial to trial.

Equations for the one-step update of the belief estimates and related HGF quantities, as well as the alternative response models, are provided in the *Supplementary Materials*.

### SimNIBS

The electric field distribution induced by each tDCS condition was simulated in each participant with the freely available SimNIBS 2.1 software [30, 31]. SimNIBS integrates different tools, such as FreeSurfer, FMRIB’s FSL, MeshFix, and Gmsh [56]. Using the headreco head modeling pipeline of SimNIBS, the electrically most relevant tissue structures (skin, skull, cerebrospinal fluid, gray matter, white matter, eyes, and air) were first segmented from the individual T1-weighted anatomical MRI. The segmentation image of the skin tissue was subsequently smoothed to remove any residual artifact. This was carried out independently from the SimNIBS pipeline with the freely available software MIPAV by applying a spatial Gaussian filter (2 mm in each xyz direction). The creation of the head model was then completed with headreco by generating a tetrahedral mesh as volume conductor model. Next, in the SimNIBS GUI, simulated electrodes were placed manually on the head mesh at their precise position and with the corresponding orientation. Stimulation intensities were selected for anodal and cathodal electrodes and the simulation based on the finite element method (FEM) was initiated. The vector norm of the electric field (normE) was extracted and chosen as dependent variable for subsequent group-level statistical analysis. These steps were repeated separately in each active tDCS condition and in each participant.

The individual normE distribution was transformed to the fsaverage space to create a group average of the mean normE values and their standard deviation (**Figure S2**). This was carried out with the MATLAB scripts provided in the SimNIBS package. To assess statistical differences between stimulation conditions in the peak values and focality of the induced electric field, we extracted the 99.9 percentile value of the normE distribution, as well as the volume in which the normE values reached the 99.9 strength percentile.

### Statistical analysis

Statistical analysis was performed with the use of non-parametric permutation tests. Pair-wise comparisons between active and sham stimulation conditions were evaluated using pair-wise permutation tests for matched samples. Factorial analyses with factor Block (1-3 levels) and Stimulation (active, sham) were carried out using synchronised rearrangements [34]. Effects were considered significant if (*P* < 0.05). In the case of multiple comparisons, we controlled the false discovery rate (FDR) at level q = 0.05 by means of an adaptive two-stage linear step-up procedure [57]. Significant effects after FDR-control are reported as (*P* < *P*_*FDR*_). Non-parametric effect sizes and corresponding confidence intervals are also provided using the probability of superiority for dependent samples Δ_*dep*_, range 0-1 [35].

